# Modulation of input sensitivity and output gain by retinal amacrine cells

**DOI:** 10.1101/273730

**Authors:** Neda Nategh, Mihai Manu, Stephen A. Baccus

## Abstract

The prevailing hierarchical view of the visual system consists of parallel circuits that begin in the retina, which then sum effects across sequential levels, increasing in complexity. Yet a separate type of interaction, whereby one visual pattern changes the influence of another, known as modulation, has received much less attention in terms of its circuit mechanisms. Retinal amacrine cells are a diverse class of inhibitory interneurons that are thought to have modulatory effects, but we lack a general understanding of their functional types. Using dynamic causal experiments in the salamander retina perturbing amacrine cells along with an unsupervised computational framework, we find that amacrine cell modulatory effects cluster into two distinct types. One type controls ganglion cell sensitivity to individual visual features, and a second type controls the ganglion cell’s output gain, acting to gate all features. These results establish three separate general roles of amacrine cells – to generate primary visual features, to use context to select specific visual features and to gate retinal output.

## Introduction

Visual computations arise from the combination of basic elements known as visual features, which are stimuli that match specific spatiotemporal patterns selected by neural circuitry. These visual features are combined, subtracted and divided to generate more complex computations. The classical or linear receptive field of a neuron is typically considered to represent the primary visual feature about which the neuron communicates. Input to the linear receptive field is often modulated by other visual features that represent a context^1^. Context-dependent modulatory effects can be spatiotemporal including the specific computations of object motion sensitivity^2^, peripheral excitation^3,4^ and surround suppression in retina^5,6^ and cortex^7,8^. The context can also be purely temporal, with the previous history defining the response to a more recent stimulus, as in the case of the omitted stimulus response^9^ and retinal sensitization^10^. Context-dependent modulation also plays a more general statistical role in visual processing, including divisive normalization, which has been proposed to reduce statistical dependencies between visual features^11^ and in non-sensory systems^12^.

Mechanisms of context-dependent effects in the retina have focused on amacrine cells, a class of diverse inhibitory interneurons that participate in multiple forms of inhibitory interactions including feedback, feedforward and lateral inhibition, targeting bipolar cells, ganglion cells, and other amacrine cells^13–15^. It has been proposed based on modeling of ganglion cell visual responses and pharmacology that amacrine cells could modulate the gain of ganglion cell visual responses^16,17^. Inhibition can reduce the gain of bipolar cells presynaptically^18^, and amacrine cells can modulate direction selective ganglion cells^19^, but how specific amacrine cells exert these influences on specific ganglion cell visual features has not been directly measured. As such, the computational role of amacrine cells in context-dependent modulation is not clear.

Here we take a general approach to directly measure context-dependent modulation driven by an interneuron, defined quantitively as a visual feature that represents the context that changes in the slope of the response to an orthogonal visual feature. This definition allows us to cleanly separate amacrine contributions that create the ganglion cell linear receptive field from modulatory effects. We focus on purely temporal aspects of processing as a first step to understand general mechanisms of amacrine cell modulation.

We used a causal manipulation approach to record from and inject current into single amacrine cells while simultaneously recording from multiple ganglion cells in the isolated retina of the salamander^20–24^. We then devised a computational framework to decipher the effect of individual amacrine cells on multiple visual features represented by a ganglion cell. We found a range of effects from amacrine cells that include both mediating a component of the average visual feature encoded by ganglion cell, and diverse modulatory effects on the individual constituent visual features conveyed by other neural pathways to the ganglion cell. The range of modulatory effects on the ganglion response to visual features affected a number of different properties, including divisive gain control, shifting of threshold, changing of sensitivity, an additive inhibitory effect, and reversal of polarity. Furthermore, even amacrine cells with simple, linear responses under stimulation by a uniform visual field could create linear and diverse types of nonlinear effects on multiple different features of single target ganglion cells.

We found unexpectedly that amacrine cell nonlinear modulatory effects clustered into two types, the first was a change in sensitivity and threshold that was specific to single visual features sensed by a ganglion cell, and the second was a change in the response amplitude that acted more uniformly on all features of a ganglion cell. These results show that an amacrine cell’s receptive field defines a modulatory context, and that stimuli matching this context drive two types of modulation – selecting among a ganglion cell’s input visual features, and exerting independent control over the saliency of the ganglion cell’s output. As context dependent modulation in the form of input-specific effects and control of output gain has also been observed in the visual cortex^25^ and auditory cortex^26^, our results give a quantitative definition and causal determination of a mechanism for a generalized sensory computation.

## Results

### The visual stimulus feature transmitted by an amacrine pathway

When considering the sensory contribution of an interneuron, one can conceptually divide signals into two broad components, those flowing through the interneuron, and those traveling to the output cell through pathways parallel to the interneuron. The interaction between those two pathways has been divided into two classes, a linear or classical effect^27–32^, whereby an interneuron contributes to the linear or classical receptive field, and a modulatory effect whereby an interneuron interacts nonlinearly with the output of other parallel pathways^2,33–41^.

To directly measure these two classes of effects from amacrine cells, we intracellularly measured (in control experiments) and then manipulated (in current stimulation experiments) the activity of individual Off-type amacrine cells including sustained (n = 8) and transient (n = 3) in the isolated salamander retina and simultaneously recorded the spiking activity of many ganglion cells (n = 153) in response to visual stimulation and amacrine perturbation using an array of extracellular electrodes (Fig. 1; see Methods).

**Figure 1.**
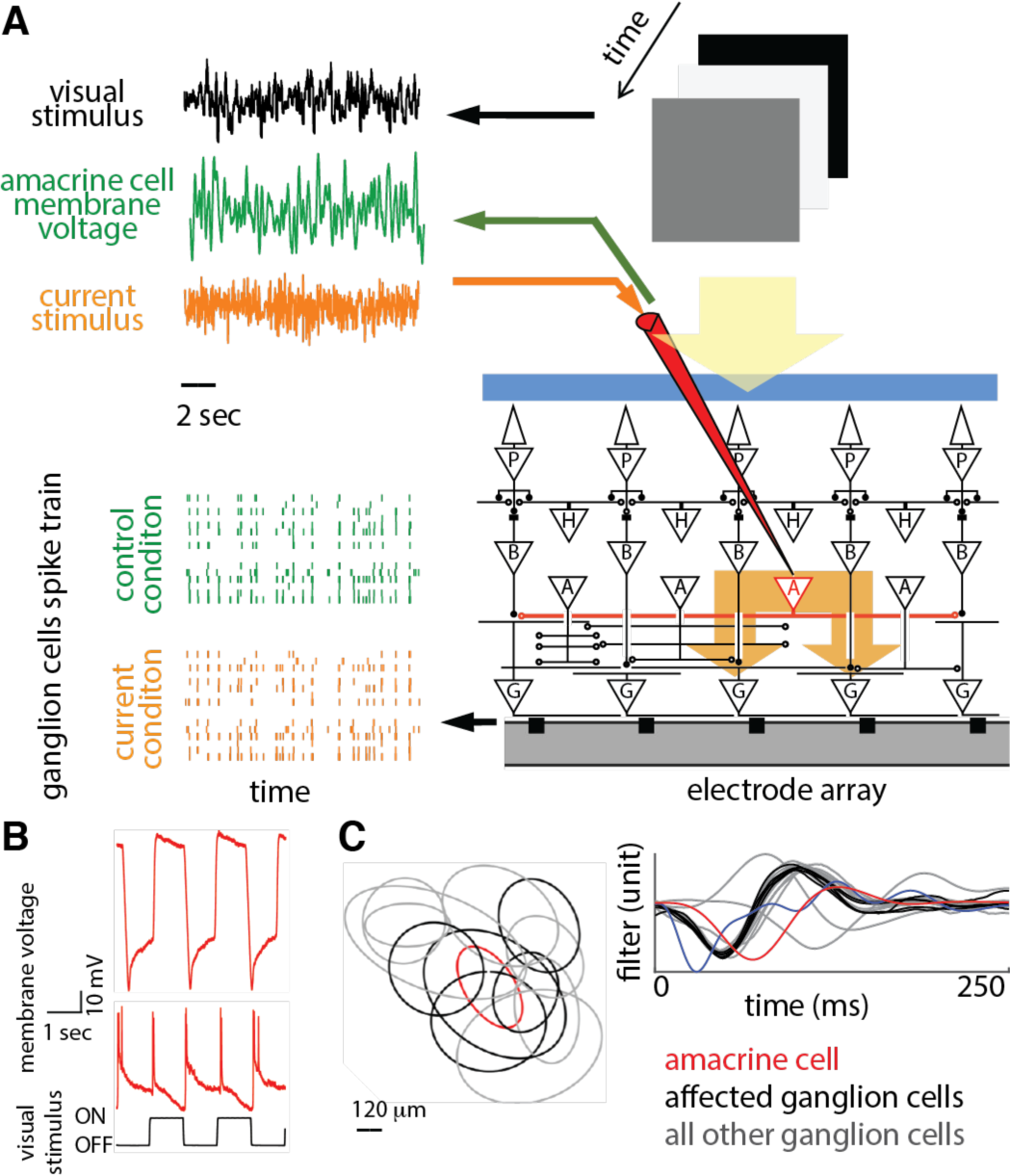
Measuring responses and transmission in an amacrine cell pathway. (A) Diagram of experimental setup for simultaneous intracellular and multielectrode array recording: a uniform field randomly flickering visual stimulus with a white noise Gaussian distribution is projected through a video monitor onto an intact isolated salamander retina. Simultaneously, Gaussian white noise current (orange trace) was injected intracellularly into an amacrine cell (current condition) or the amacrine cell membrane voltage (green trace) was recorded intracellularly using a sharp microelectrode (control condition). A multielectrode array of 60 extracellular electrodes recorded spiking activities of multiple ganglion cells simultaneously. Multiple traces on the left represent spiking response of multiple ganglion cells to a single stimulus trial during the control condition (orange spikes) and current condition (green spikes). (B) The membrane potential response of a sustained Off-type amacrine cell (top row), the response of a transient On–Off type amacrine cell (middle row), to a uniform field flashing stimulus (bottom row). (C) (Top) Receptive fields of a single amacrine cell and multiple ganglion cells. Each oval indicates one standard deviation of a two-dimensional Gaussian fit to the receptive field mapped using a white noise checkerboard stimulus. Red oval indicates an amacrine cell receptive field; black ovals indicate ganglion cells affected by current injection into the amacrine cell. Grey ovals are unaffected ganglion cells. (Bottom) Temporal filters computed for recorded cells, including: amacrine visual filter (red), computed by correlating a uniform-field visual stimulus and the amacrine cell membrane potential; ganglion cell visual filters, computed by correlating the visual stimulus and ganglion spikes for the affected ganglion cells (black) and all other recorded ganglion cells (grey); and an amacrine transmission filter, between the amacrine cell and one affected ganglion cell (blue) computed by correlating a white noise current stimulus injected into the amacrine cell and the ganglion cell’s spikes.

We first analyzed the visual feature transmitted by the amacrine cell, which is formed in two stages – the amacrine cell’s visual response, and its transmission. We measured the visual response of amacrine cells by fitting a linear-nonlinear (LN) model consisting of a linear temporal filter followed by a time-independent, or static nonlinearity (Fig. S1A). The linear filter represents the average effect of a brief flash of light on the amacrine cell membrane potential, or equivalently can be interpreted as the time-reverse of the visual stimulus to which the amacrine cell was most sensitive on average.

To characterize the second stage of the interneuron pathway – how signals transmitted through individual amacrine cells contributed to the ganglion cell response – we injected white noise current into the amacrine cell while presenting a white noise visual stimulus. Because the inner retina adapts to the stimulus contrast by changing its sensitivity^42,43^, the visual stimulus was included to maintain the retina in a similar state of adaptation in the current injection and control conditions. We then computed an LN model between the amacrine current and the ganglion cell spike train (see Methods, Fig. S1B). In this transmission model, the transmission filter represents the average effect of a brief pulse of current on the ganglion cell’s firing rate. The linear transmission filters were mostly inhibitory with different time courses (Fig. S1C). The efficacy of transmission between each pair of amacrine cell and ganglion cell was assessed based on a significant peak in the measured linear transmission filter as compared with a control filter computed by shuffling the ganglion cell’s spike train in time. Thirty-nine out of 153 measured transmission filters met this criterion (Fig. 1C).

Because the input signal to this cascade has a Gaussian probability distribution, the overall linear feature conveyed by this pathway, referred to as the amacrine pathway filter, can be found by convolving the two linear filters whether the amacrine visual response is linear or nonlinear^44^. In combining these two filters, we avoided a double contribution from the membrane time constant by deconvolving an exponential filter representing that time constant^22^. The features conveyed by the amacrine pathway were typically monophasic and positive, consistent with previous measurements^45^, formed by a negative response to light (Off response) followed by an inhibitory transmission; there was diversity in the time course of the amacrine pathway features (Fig. S1D).

Effects were seen on only a subset of ganglion cells (Fig. S1E-F). Consistent with previous results from steady current pulses^21^, transmission dynamics in some cases were specific to the amacrine–ganglion cell pair, with single amacrine cells showing different dynamics for different target ganglion cells (Fig. S2). In the following, the analysis of amacrine cells’ effect on the downstream ganglion cells is performed only on the amacrine–ganglion cell pairs for which a significant transmission filter could be measured.

### Signals traveling through parallel neural pathways

To identify the visual input conveyed to a ganglion cell by neural pathways other than through the causally perturbed amacrine cell, we define the visual stimulus space that is the complement to the amacrine pathway visual feature that was captured by the amacrine pathway filter described above. To do this, we removed the amacrine pathway contribution out of the *n*-dimensional visual stimulus by computing the (*n-1*) dimensional stimulus space orthogonal to the amacrine pathway feature (see Methods, and ^46,47^). Within this new stimulus subspace, which represents the space of features within which the amacrine cell might exert modulatory effects, we computed the ganglion cell’s spike-triggered average (STA) feature, the orthogonal STA (oSTA) (Fig. 2A). This feature represents the average of other visual features encoded by the ganglion cell, that were not conveyed by (were orthogonal to) the amacrine pathway. This two-pathway model thus encodes two properties of the stimulus, one average feature that we causally measure is encoded by the recorded amacrine cell, and one average feature that is encoded by all other interneuron types. We confirmed that this two-pathway model–using the amacrine transmission filter measured in the ‘current stimulation’ experiments, more accurately predicted ganglion cell responses than the LN model when applied to the ganglion cell’s spiking responses recorded during the ‘control experiments’. The correlation coefficient between model and data for the two-pathway model was greater than that for the LN model (0.10 ± 0.02 mean ± SEM, p = 1.65 x 10^-5^, Wilcoxon signed-rank test, n = 39) on held-out data not used for estimating the model (Fig. S3).

**Figure 2.**
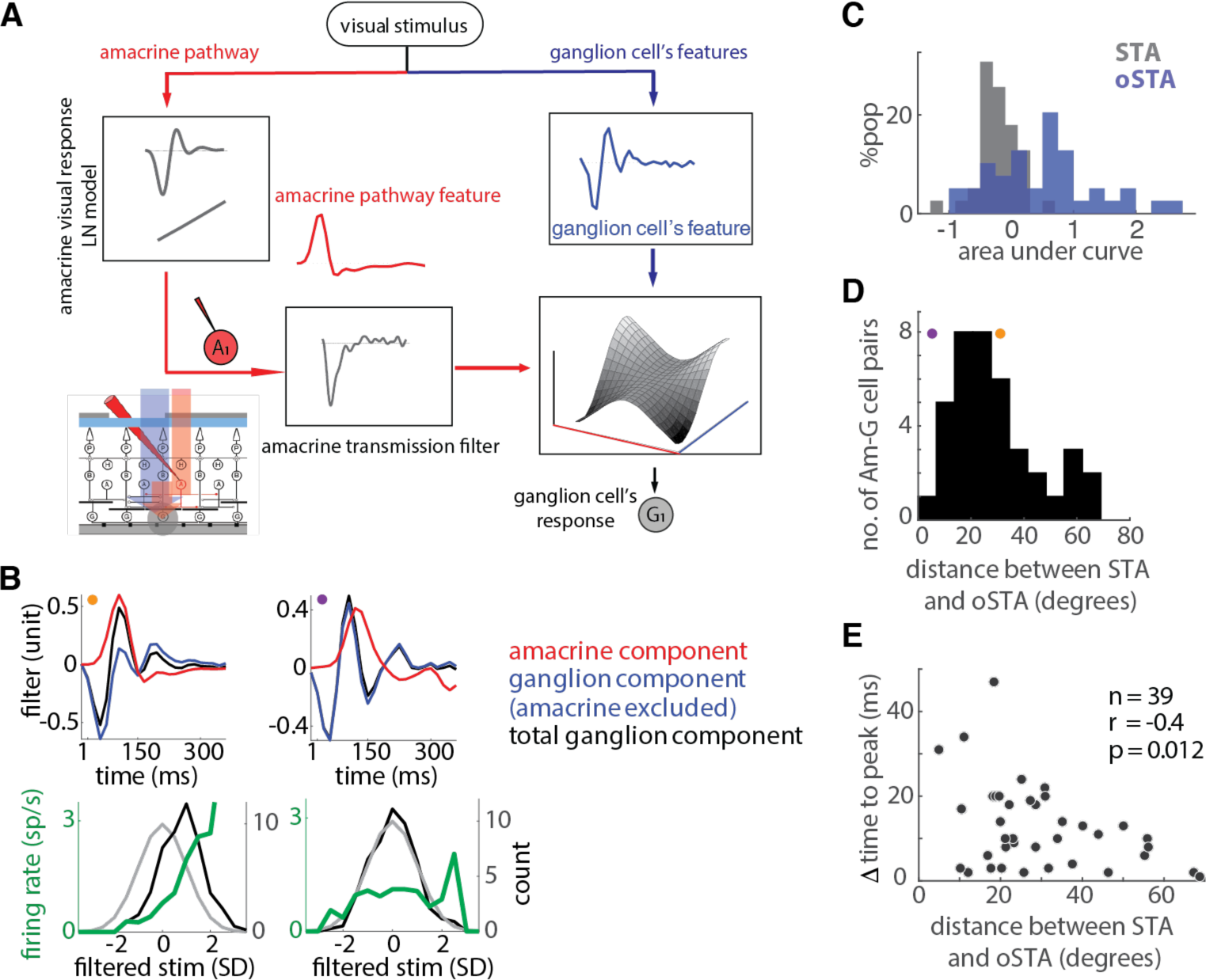
Contributions of amacrine cells to the average ganglion cell visual feature. (A) Illustration of a model containing two main pathways, an amacrine pathway whose feature is shown by the red linear filter, and another pathway representing other ganglion cell features whose average is shown by the blue linear filter. The outputs of these two pathways are combined by a two-dimensional nonlinear function to generate the ganglion cell’s firing rate. (B) (Left column, top) Sample amacrine–ganglion cell pair for which amacrine pathway feature (red curve) contributed to the ganglion cell’s STA (black curve) as indicated by the observation that the STA and the oSTA (blue curve) are different. (Left column, bottom) The raw stimulus distribution (light grey) and the spike-triggered stimulus ensemble distribution (black), as well as the amacrine pathway nonlinear response function (computed as a quotient of the spike-triggered and raw stimulus distributions) (green), when the stimulus is projected onto the amacrine pathway feature (red). (Right column) Same as left column for an amacrine cell for which the oSTA was similar to the STA. (C) Histogram of center-surround weighting for STA (black) and oSTA (blue) across 39 amacrine–ganglion cell pairs, expressed as the signed area curve of their STA and oSTA filters. (D) Histogram of the observed difference between the STA and oSTA for 39 amacrine–ganglion cell pairs, expressed as the angle difference in degrees. Colored symbols indicate the angle differences corresponding to the sample cell pairs shown in (B). (E) Relationship between difference between the time to peak of an amacrine pathway filter and the target ganglion cell STA (y-axis) versus the difference between STA and oSTA across 39 amacrine–ganglion cell pairs.

The oSTA is the average feature of a subspace of the stimulus orthogonal to the amacrine contribution. Figure 2B shows the measured STA and oSTA for two example amacrine–ganglion cell pairs. The population of ganglion cell STAs had a very similar center-surround weighting (area under curve 0.22 ± 0.23 median ± MAD (mean absolute deviation)), consistent with previous results^45^. The oSTA was more diverse (area under curve -0.57 ± 0.65 median ± MAD; p < 0.001, paired ttest between MAD of bootstrapped STA and oSTA area under curve distributions), which likely arose because there is a range of the strength of the amacrine cell to the ganglion cell linear receptive field as previously described (Fig. 2C).

For some cells, the oSTA was different from the STA (Fig. 2B, left), whereas for other cell pairs the oSTA and STA were highly similar (Fig. 2B, right), meaning that the amacrine cell’s contributed feature was orthogonal to the STA. On average, the distance between ganglion cell’s STA and oSTA was 25.14 ± 12.61 degrees median ± MAD across 39 amacrine–ganglion cell pairs (Fig. 2D). The similarity of STA and oSTA was negatively correlated with the timing difference between the peaks of the amacrine feature and the STA, such that when the amacrine cell’s feature and the ganglion cell STA had similar peaks, the STA and the oSTA were more different (Pearson correlation: -0.4, p = 0.012) (Fig. 2E). Figure S4 provides more examples of the amacrine cells’ contribution that were used for the population analysis in figure 2D.

### A general analysis of modulation in amacrine pathways

A ganglion cell receives input from multiple neural pathways, and each pathway conveys its own preferred stimulus feature. An interneuron could potentially affect each of these features differently. We were interested in understanding (1) what ganglion cell visual features are affected by the amacrine pathway? (2) what kind of nonlinear interaction exists between the amacrine pathway and other input pathways to a ganglion cell? and (3) are the amacrine modulatory effects different for different features? To address these questions, we examined further the stimulus space orthogonal to the amacrine preferred feature. The oSTA is the average of this orthogonal stimulus ensemble, but to analyze modulatory effects of the amacrine cell we decomposed this stimulus ensemble into its principal components, each representing a separate feature, with all of these features being orthogonal to the amacrine feature. We used the standard approach of spike triggered covariance (STC) analysis to recover this entire linear subspace orthogonal to the amacrine feature, representing the space spanned by the features of the multiple pathways feeding into the ganglion cell, and that is not linearly encoded by the amacrine pathway. We define this subspace as the modulated subspace of the amacrine cell. Using this orthogonal STC approach, for each ganglion cell, we recovered up to several modulated features that were orthogonal to the amacrine feature (Fig. 3A, Fig. S5-S6). Note that these features do not necessarily correspond to other specific neural pathways.

**Figure 3.**
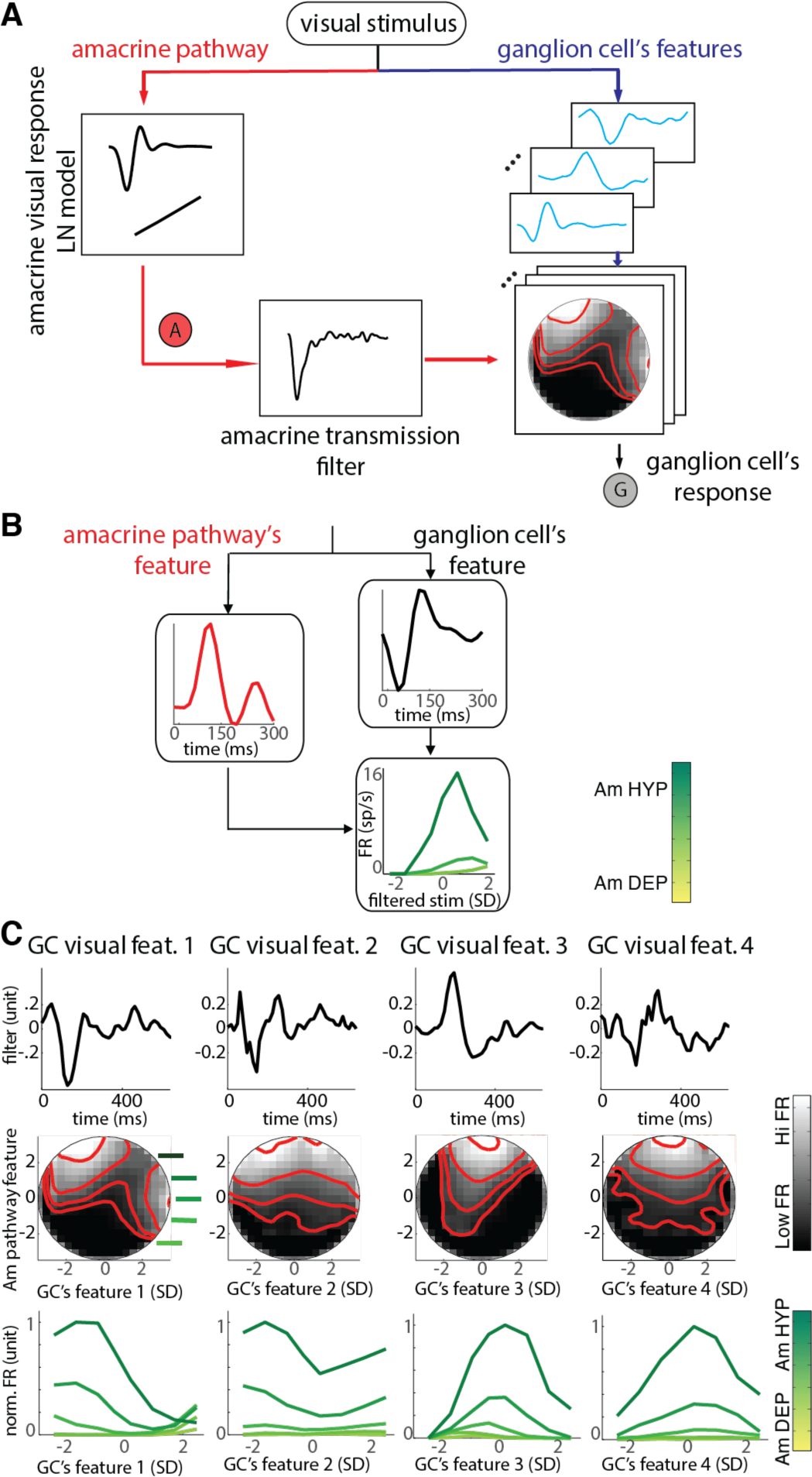
Differential modulation of multiple distinct ganglion cell visual features by an amacrine cell. (A) The multi-pathway model: The space of visual features encoded by a ganglion cell was decomposed into its principal components (light blue traces) using STC analysis on the stimulus space when the amacrine pathway feature is projected out. Then for each of the ganglion cell’s feature dimensions and the amacrine pathway’s feature dimension, a two-dimensional nonlinear firing rate function is computed. (B) (Left) Illustrates a nonlinear interaction between the amacrine pathway and one of the orthogonal STC features in (A) for a pair of an amacrine cell and a ganglion cell. Red trace represents the amacrine pathway feature; black trace represents the orthogonal STC feature; and green traces show one-dimensional nonlinear response functions computed from the two-dimensional instantaneous firing rate nonlinearity for four bins of amacrine pathway output values specified by the colorbar. (C) Amacrine–ganglion cell’s features nonlinearity characterization: (Top) Orthogonal STC significant dimensions representing the ganglion cell’s other features, excluding the amacrine pathway feature, for a sample amacrine–ganglion cell pair. (Middle) Two-dimensional firing rate nonlinearity as a function of the amacrine pathway output (y- axis) and the projection of the stimulus on each orthogonal feature (x-axis). The y-axis is the projection of stimuli on the amacrine pathway feature, and the x-axis is the projection of the stimuli on the corresponding ganglion cell’s visual feature. The two-dimensional instantaneous firing rate was computed as a quotient of the spike-conditional and raw stimulus distributions. Lighter regions correspond to higher ganglion cell firing rate (FR) in the two-dimensional stimulus subspace. (Bottom) One-dimensional slices of each two-dimensional firing rate as a function of the projected stimulus onto the corresponding ganglion cell’s feature (x-axis) for different levels of the amacrine cell’s feature output. Trace color indicates different levels of the amacrine feature output, indicated by green bars in the middle row. Because the amacrine cell is inhibitory (its transmission filter has a negative peak), a high level of the amacrine pathway’s preferred feature roughly corresponds to the amacrine cell being more hyperpolarized (darker green colors).

To quantify how the amacrine feature modulated each of the ganglion cell features, we computed for each modulated feature a two-dimensional nonlinear function that combined the amacrine output and the modulated feature. The inputs to this 2D nonlinearity were the projection of the stimulus on the amacrine pathway feature, and the projection of the stimulus on the modulated ganglion cell feature (Fig. 3A). The output of the 2D nonlinearity was the ganglion cell firing rate as a function of the similarity of the stimulus and the modulated ganglion cell visual feature, for different levels of the amacrine pathway output (Fig. 3B).

Several different categories of effects were observed in these 2D nonlinearities. Figure 3C shows the ganglion cell’s modulated features (Fig. 3C top row) for an example amacrine–ganglion cell pair, and the ganglion cell’s 2D response nonlinearity as a function of the stimulus projection onto the amacrine pathway feature and each of the modulated ganglion cell’s features (Fig. 3C middle row). Figure 3C bottom row shows a different view of these 2D nonlinearities as a series of input-output functions (ganglion cell modulated feature input and firing rate output), i.e., 1D nonlinear functions that change (are modulated) depending on the amacrine output. For different modulated visual features of the same amacrine–ganglion cell pair, this view revealed different qualitative types of amacrine effects on different ganglion cell features. These included 1) an additive effect on firing, such that the baseline level was high when the amacrine cell was hyperpolarized, 2) a change in sensitivity such that visual input that matched the visual feature caused a greater change in ganglion cell firing when the amacrine cell was hyperpolarized, 3) suppression of the ganglion cell visual feature by amacrine depolarization, either with or without a change in threshold to visual input. Some of the ganglion cell visual features exhibited nonmonotonic nonlinearities, possibly reflecting a combination of multiple pathways connecting to a ganglion cell. In addition, more complex effects were observed, including that as the amacrine cell changed from depolarized to hyperpolarized, the ganglion cell response changed from Off– type to On–type (first column in Fig. 3C), consistent with previously reported polarity reversal during visual stimulation^48^. Our analysis shows that a local amacrine pathway can change the sign of a ganglion cell’s response to light on a fast time scale. Overall, these results illustrate how even under a constant contrast white noise stimulus, inhibitory interneurons can rapidly and dramatically alter sensitivity to visual features in the course of a computation.

Note that even though the amacrine effect on the overall firing rate is inhibitory for all of these features, in the sense that whenever amacrine cell is hyperpolarized there is greater firing, the nature of the interaction of the amacrine output with the ganglion cell preferred feature is very different. This analysis reveals that simply designating a cell as inhibitory is insufficient to account for the complexity of its effects. Furthermore, even in a single amacrine–ganglion cell pair multiple types of effects can occur, which suggests multiplexing at the level of a single inhibitory neuron. The diversity in nonlinear modulatory effects exists both for a single amacrine cell across different visual features of a single ganglion cell and for a single amacrine cell across multiple target ganglion cells (Fig. S7). Taken together, using white noise visual and current stimulation combined with a subspace cascade model approach enabled us to develop an unbiased methodology for dissecting the modulatory effects of an interneuron. Our approach yields a quantitative and precise definition of modulation – an interneuron’s linear feature defines the context, and that context modulates orthogonal visual features.

We could identify five classes for the nonlinear modulatory effects of the amacrine pathway on the ganglion cell’s light response, which included changing the sensitivity, controlling the gain, changing the threshold, an additive inhibitory effect, and reversal of polarity, or alternating between Off– and On–type ganglion cell (Fig. 4). These classes do not represent separable types, but axes defining different effects. Figure 4A shows the computed response function for example amacrine– ganglion cell’s feature pairs in our dataset, demonstrating the five categories of the amacrine pathway’s modulatory effects. The effects found for the pairs studied here were not specific for a particular amacrine cell or ganglion cell type, but were specific for each amacrine–ganglion cell’s feature pair. Figure 4B shows how these categories correspond to a shifting and scaling of the one-dimensional nonlinear functions across different polarizations of the amacrine cell.

**Figure 4.**
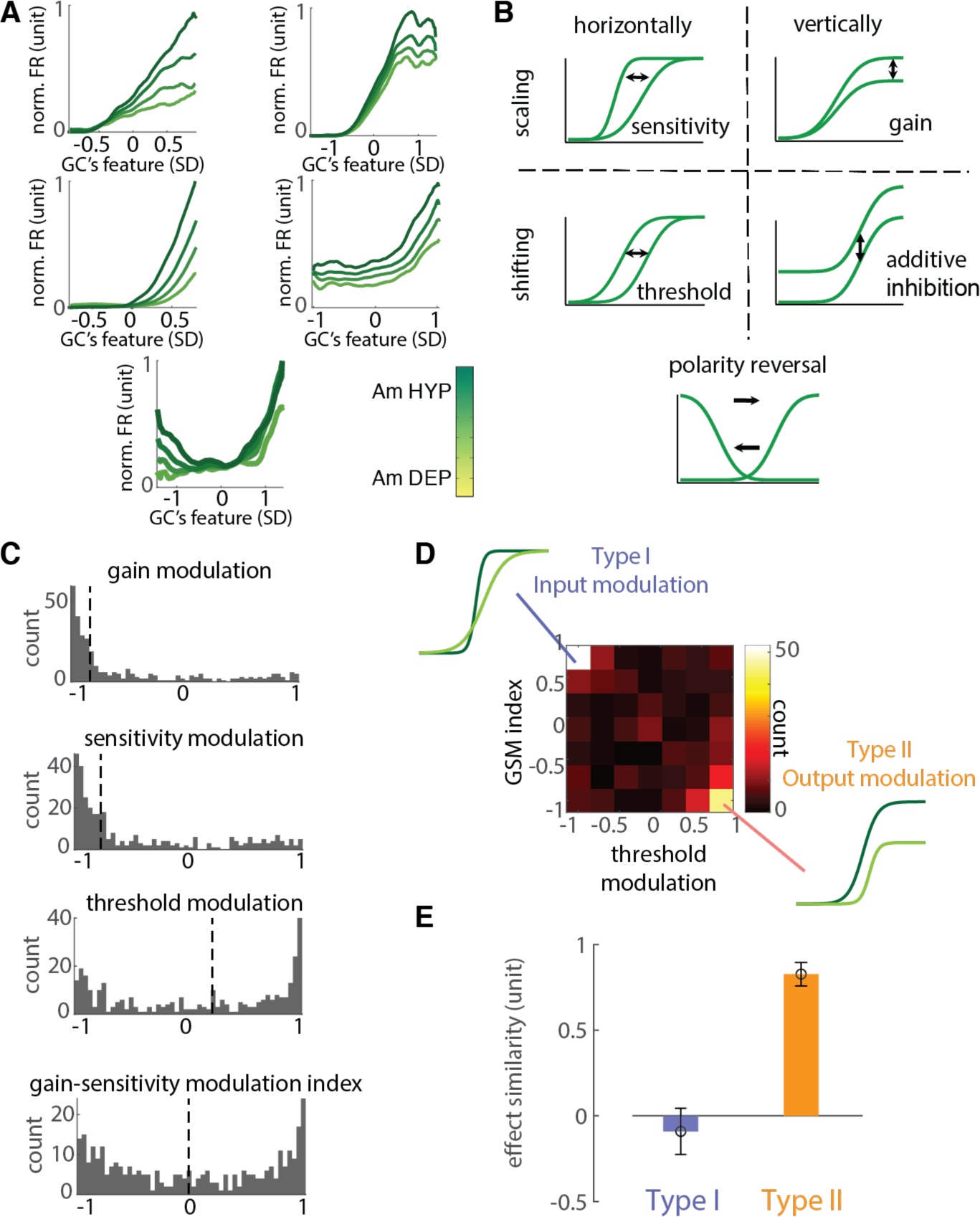
Two types of modulatory effects of amacrine cells on the ganglion cell population. (A) Firing rate response nonlinearities from example amacrine–ganglion cell’s feature pairs, representing five different types of nonlinear modulatory effects observed across 321 amacrine– ganglion cell’s feature pairs. The amacrine cell’s nonlinear effects include combinations of additive and multiplicative transformations of the ganglion cell’s nonlinear response function as the output of the amacrine pathway varied from weaker to stronger inhibition. (B) Illustrations of how changes in threshold, sensitivity, gain, or offset and polarity reversal as shown for sample amacrine– ganglion cell’s feature pairs in (A) can be obtained by simple additive or multiplicative operations on the ganglion cell’s nonlinear response function. (C) Distribution of different types of nonlinear modulatory effects identified by a multi-pathway model framework across 321 amacrine–ganglion cell’s feature pairs, which include changes in the response gain, sensitivity, threshold, and gain- sensitivity modulation (GSM) index, estimated via fitting a piecewise linear approximation of a sigmoidal function to the ganglion cell’s nonlinear response function. Dashed lines indicate the median of each histogram. (D) Center heatmap shows the joint distribution of response threshold modulation and gain-sensitivity modulation index, showing how these response variables change together when inhibition from the amacrine pathway gets stronger. Each data point (n = 321) represents the Pearson correlation coefficient between different levels of amacrine pathway polarization and the corresponding values of each response variable. Inset nonlinear functions illustrate changes in the nonlinear properties of a ganglion cell’s response to distinct visual features, associated with the two types of functional effects of amacrine cells, sensitivity modulation (top left), and gain modulation (bottom right). (E) The correlation coefficient of the change in the response nonlinearity parameters across visual features of each ganglion cell for each type of modulatory effect in (D) (median±SEM), showing variation of the effect across different modulated features for the same amacrine–ganglion cell pair.

### Two types of modulatory effects on the ganglion cell population

The dynamic perturbations to single amacrine cells during visual stimuli allowed us to directly measure and analyze diverse modulatory effects on the ganglion cell population. To further classify these effects across the population of amacrine-ganglion cell pairs, we parameterized the 2D nonlinearity for each amacrine–ganglion cell’s feature pair with a different sigmoidal function at each level of amacrine output. Because this 2D nonlinearity offers a snapshot of the changing relationship between a specific visual feature and ganglion cell firing, at different levels of amacrine output, analysis of the changing parameters of this sigmoid allowed a compact definition of the fast modulatory effects for each amacrine-ganglion cell pair.

For the ganglion cell’s response at different levels of amacrine polarization, we fit a function with only four parameters to different one-dimensional response nonlinearities corresponding to each of these levels and analyzed how the amacrine cell changes these parameters. We fit the ganglion cell nonlinearity corresponding to each level of amacrine output to a piecewise linear approximation of a sigmoid function with parameters for the minimum, maximum, horizontal offset and slope. Parameters of this nonlinear model were optimized using the nonlinear least squares method with additional lower and upper bound values constraints for different parameters (for details refer to Methods).

Characterizing the ganglion cell’s response function in terms of the parameters of the response nonlinearity enabled us to trace the changes in the response characteristics as a function of amacrine pathway output. Figure 4C shows the distribution of amacrine effects on ganglion cell nonlinearities. We computed the modulation of response sensitivity, gain, and threshold as a function of amacrine pathway output for 321 pairs of amacrine and ganglion cell features where the ganglion cell response function met the goodness of fit criteria for the nonlinearity. In addition, to aid in classifying different effect types, we computed a gain-sensitivity modulation (GSM) index, defined as the correlation between amacrine output and the ratio of response gain to sensitivity. This index represents the extent that amacrine output changes ganglion cell response vertical scaling compared to horizontal scaling. As shown in figure 4C, the threshold modulation and gain- sensitivity modulation index showed a bimodal distribution, whereas a skewed distribution was found for modulation of gain and sensitivity.

When we examined the joint distribution of gain-sensitivity modulation index and threshold modulation, we observed that there were two clusters of effects (Fig. 4D, Fig. S8). During strong amacrine inhibition, one effect (Type I) showed a positive GSM index (associated with greater decrease in sensitivity) along with a significant decrease in threshold, whereas another effect (Type II) showed a negative GSM index (associated with greater decrease in gain) along with a significant increase in threshold. In addition, there was overall a strong negative correlation between the GSM index and change in threshold (Pearson correlation: -0.55, p < 10^-26^).

Within each of these two clusters, amacrine inhibition either affected gain or sensitivity, but largely not both. For the Type I effect, the GSM index was correlated with a change in sensitivity but not with gain (r _GSM index-gain_ = 0.01±0.07, r _GSM index-sensitivity_ = -0.60±0.05; mean±SEM, Pearson correlation) (Fig. 4D, top left region). For the Type II effect, the modulation index was much more strongly correlated with a change in gain than with sensitivity (r _GSM index-gain_ = 0.57±0.05, r _GSM index- sensitivity_ = -0.27±0.06; mean±SE, Pearson correlation) (Fig. 4D, bottom right region). Overall, these results indicate primarily two types of effects between amacrine and ganglion cell pairs, one that more strongly influenced sensitivity, and one that affected gain, with both effects occurring with a shift of threshold.

We further analyzed whether modulatory control over different visual features might act separately on specific visual features or act more similarly on all features together. For each single amacrine cell-ganglion cell pair, we analyzed the similarity of effects on different affected visual features by computing the correlation coefficient across visual features of the change in parameters of sensitivity, gain and threshold for the sigmoidal visual response curve (Fig. 4E; see Methods). We found that for Type I effects, the change of parameters for different visual features was uncorrelated (r = -0.09±0.14_median±SEM, Pearson correlation, 19 cell pairs; p = 0.79 Wilcoxon signed-rank test against zero correlation), indicating that amacrine modulation was feature-specific – effects on different visual features were different. However, for Type II effects, the change in parameters were significantly more correlated across modulated visual features for the same amacrine-ganglion cell pair (r = 0.83±0.07_median±SEM, Pearson correlation, 13 cell pairs; p = 0.006 one-sided Wilcoxon rank-sum test). This type of modulation acted more to control the gain of visual features together, consistent with modulation of the ganglion cell’s output. Thus overall, we quantitatively characterized two types of modulatory effects caused by an amacrine cell (Fig. 5), one of input modulation that acts primarily to control the sensitivity of visual features in a feature-specific manner, and one of output modulation that acts more to control the gain of visual features of a ganglion cell together.

**Figure 5.**
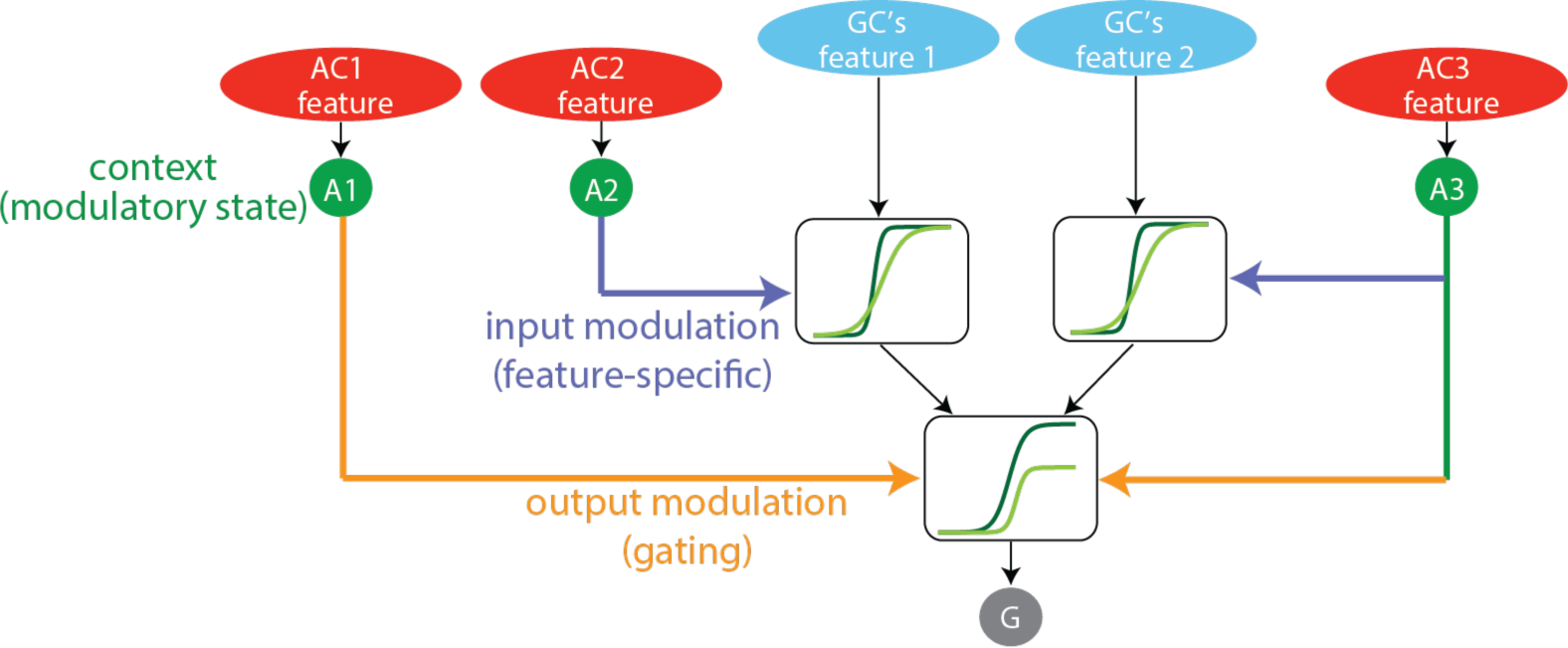
Amacrine cells create two types of context-dependent modulation of ganglion cells. Schematic diagram of how visual features of different amacrine cells represent a context that drives two types of modulation. The first is the control of input sensitivity of visual features prior to summation within the ganglion cell. The second is control of the output gain of the ganglion cell, which acts similarly on all visual features of the ganglion cell. Colors used for differentiating different connections match those used in figure 4D-F.

## Discussion

We have dissected the functional effects of an interneuron in a circuit using measurements of the interneuron’s response, dynamic causal perturbations of the interneuron, and simultaneous recording from a population of downstream cells. White noise analysis in a cascade and subspace model framework disentangled the linear and nonlinear contributions of the interneuron in an unsupervised and systematic manner. From these experiments and analyses, we have characterized different types of functional effects arising from the same amacrine cells. The first of these effects is the linear contribution in which the amacrine pathway helps to build the linear receptive field of the output neuron^45^. Such effects show that some amacrine cells contribute directly to the average ganglion cell stimulus selectivity. Secondly, by measuring the nonlinear effects of the amacrine cells, we observed modulatory effects of these inhibitory interneurons on the ganglion cell’s response to specific visual features, with different effects occurring on different features. The visual feature conveyed by the amacrine cell served as a context to drive the modulation of other, distinct visual features. At the lowest level, these effects include a shift in threshold of the ganglion cell’s response to a visual feature, the control of input sensitivity and output gain. When we examined how these component effects occurred together, we saw two different types of effect combinations that could be observed even with the same amacrine– ganglion cell pairs. Input modulation that primarily controlled sensitivity rather than gain, and acted specifically on individual visual features, and output modulation that primarily controlled gain, and acted more similarly across different visual features. These effects are modulatory, meaning that one feature – the amacrine feature – changes the response of the ganglion cell to other features. Across many amacrine cells with different effects, the amacrine cell population defines a context that modulates multiple other visual features conveyed by the ganglion cell population.

### Linking a Functional Pathway to a Neural Pathway

Of central importance in the study of sensory processing are the stimulus features that drive the neural response. Numerous studies have shown that many neurons respond to more than one feature in a high-dimensional stimulus space^49–58^. Although dimensionality reduction approaches, such as spike-triggered covariance (STC) and its extensions^59–64^ or maximally informative dimensions (MID)^65–67^, have been successful to find a relevant linear subspace that causes a change in the neuron’s response, the resulting features cannot be easily interpreted in terms of functional computations. As a result, this subspace description provides little insight into the neural circuitry that gives rise to the neuron’s response. Moreover, it is not clear that dimensions extracted by these methods relate to neural pathways, and we only know that the space of these extracted dimensions encompasses the space of features encoded by all the neural pathways feeding into one neuron. Although other work has tried to rank these stimulus dimensions according to how much information they capture^63^, it is still difficult to link the resulting features to the neural pathways encoding those features.

In this study, by directly measuring the amacrine pathway response and effects of transmission, we can assign a neural identity to those features and characterize modulatory effects in a way that would not be possible by modeling based on the ganglion cell response alone. We see for the first time using a direct, causal, general and assumption-free approach how the widely studied computation of context-dependent modulation is constructed by interneurons.

### Potential origin of modulatory types

In our study we have identified that amacrine cell inhibition provides a mechanistic basis for two types of modulation of visual features. Looking down one level further, future studies will address the cellular basis for the two different modulation types. One might think that changes in sensitivity might occur prior to the strong threshold of the bipolar cell terminal, and thus might be presynaptic, and that changes in gain might arise from inhibition delivered after this threshold postsynaptically. Fitting with this idea, the feature-specific effects of Type I modulation are consistent with a presynaptic effect on individual bipolar cells, and the more feature-wide effect of Type II is more consistent with a postsynaptic effect. However, more complex effects of presynaptic adaptation could complicate this picture, coupling effects on sensitivity with effects on threshold or gain^68^. Our study nevertheless provides a testable hypothesis that the origin input modulation (Type I) is presynaptic and the origin of output modulation (Type II) is postsynaptic. Note that in other systems, the more relevant result may be the two types of modulation, rather that the specific underlying cellular basis. For example, in the cortex, because dendritic integration can be local and occur prior to spiking in dendrites, input modulation could be delivered to dendrites, and output modulation could be delivered closer to the soma^69^.

### Implications of amacrine cell modulation for natural scene processing

It has been suggested that neural representations of sensory stimuli are optimized to reduce redundancy based on the statistics of the natural environment, thereby creating a more efficient representation^70–76^. These theories of efficient coding have derived an optimal set of linear basis functions that have properties similar to receptive fields in the visual cortex^77–79^. However, the statistics of the natural signals are too complex to be optimally decomposed by only a linear transformation. Moreover, neural responses are highly nonlinear, for example due to rectification or saturation of responses^80–87^. Recent studies in primary visual cortex V1 and in the retina have shown that nonlinear properties play an important role in decorrelating the visual image and therefore in the efficient coding of natural signals^11,88^.

One class of multidimensional nonlinear models that has linked natural scene statistics to neural response properties exhibits so-called divisive normalization^89^. The intuition behind this model is to divide the signal by its variance as predicted from a linear combination of the variance of other features to obtain a representation where different features are represented with similar variance. The divisive normalization model describes the firing rate responses of cortical neurons as the ratio between the output of a linear receptive field and the magnitude of a set of different visual features. Such models are consistent with responses found across cortical areas and sensory modalities^90^. If different visual features often occur together, in other words they are redundant, then if one feature suppresses the other it will tend to create a less redundant representation.

Of the two nonlinear effects we observed, these effects are consistent with a key property of divisive normalization, namely that visual features reduce the sensitivity to other visual features. Our findings suggest that the normalizing directions are in fact the features encoded by the amacrine cells, and that the amacrine cell’s own preferred or optimal feature can be different from the preferred or optimal feature of the ganglion cell. We propose that a major function of amacrine cell circuits may be to undo the receptive field dependencies of multiple visual pathways by controlling the gain and sensitivity along the features encoded by each of these pathways.

### Future studies

To test this redundancy reduction hypothesis, it will be necessary to measure amacrine nonlinear modulations when the features of amacrine pathway and other ganglion cell’s incoming pathways are coincident. Therefore, it will be necessary to repeat our experimental and computational procedure while presenting a spatiotemporal visual stimulus containing the type of structure observed in natural scenes. In our current analysis, because stimuli were composed of a uniform field, we only measured the contribution of the temporal component of the stimulus. However, all amacrine and ganglion cells were within < 200 µm, and thus are highly likely to have overlapping spatial receptive fields. It is thus likely that amacrine and ganglion cell features will be activated together by stimuli with sharp transitions such as an edge. Nevertheless, it will be important to directly measure the spatiotemporal filters of amacrine and ganglion cell features. Despite the limitations of temporal stimuli that we have used, there is sufficient complexity and generality in temporal processing^10,68,91^ to support our fundamental conclusions about amacrine modulation. Furthermore, our approach can be extended to spatial stimuli in a straightforward way.

For application to other neural circuits, the techniques used in this study only require an intracellular recording and stimulation and an extracellular recording from an affected cell. Nothing in our recordings, analysis or conclusions require that amacrine and ganglion cells are monosynaptically connected, merely that one functionally affects the other. Newer technologies make it feasible for our approach to be used elsewhere in the nervous system using multielectrode recording to record affected cells. Opto-tagging could allow a presynaptic cell recorded with the multielectrode probe to be identified, and then this optogenetic stimulation would allow perturbation of those cells as we have done. Context-dependent modulation is studied widely in sensory systems, and both input modulation and output modulation has been observed, either physiologically^25,26^, or based on anatomical evidence^92,93^. Our approach provides a way to identify these modulatory effects in an unbiased manner, and account for the responsible neural pathways.

Here we have presented a systematic approach to understand the functional contribution of an interneuron. Although the anatomical diversity of amacrine cells allows for the combination of diverse visual features, we propose that these effects can now be understood in terms of three basic functions of sensory inhibitory interneurons, to generate the primary feature sensed by a neuron, to create a context that acts on specific other sensory features, and to use that context to gate a neuron’s output. These effects may represent the general actions of sensory inhibitory interneurons.

## Methods

### Electrophysiology Visual Stimulation

Visual stimuli were projected onto the retina from a video monitor, at a photopic mean intensity of 8 mW/m^2^, and at a frame rate of 63 Hz. The video monitor was calibrated using a photodiode to ensure the linearity of the display. Stimuli were spatially uniform and were drawn from a white noise Gaussian distribution with a constant mean intensity. Contrast was defined as the standard deviation divided by the mean of the intensity values. Contrast was fixed throughout the experiment, and ranged 0.06–0.25 across recording sessions. The duration of stimuli was 300–1200 s. By keeping the mean light intensity and contrast constant throughout the experiment, we avoided any contributions from light or contrast adaptation^22,43^.

### Multielectrode Recording

To record the spike trains of retinal ganglion cells, the retina of the larval tiger salamander was isolated intact and was placed on a dialysis membrane attached to a plastic holder. Then the holder was lowered onto a flat array of 61 microelectrodes (Multichannel Systems) and bathed in oxygenated Ringer’s solution buffered with bicarbonate^94,95^. Extracellular electrodes were spaced 100 μm apart. Electrical signals from the array of electrodes were digitized at 10 kHz and recorded to a computer. Results included 153 recorded ganglion cells, which comprised biphasic Off-type, monophasic Off-type, biphasic on-type, and On–Off cells classified with a uniform field visual stimulus.

### Simultaneous intracellular and multielectrode recording

Simultaneous intracellular recordings were made with sharp electrodes from eleven amacrine cells with a resting membrane potential −55 ± 4 mV (mean ± SEM), including sustained (n=8) and transient (n=3) Off amacrine cells. Amacrine cells were identified by their light responses, the presence of an inhibitory surround, and their inhibitory transmission to Off-type ganglion cells with overlapping receptive field centers. The retina was held in place over the electrode array under a 100-μm layer of 0.6% agarose, covered by a dialysis membrane containing several 100-μm holes. The intracellular electrode was then guided under infrared light through a hole and the agarose layer to penetrate the retina from the photoreceptor side. Intracellular electrodes (150–250 MΩ impedance) were filled with 1–2 M potassium acetate.

### Current Stimulation

Current was injected through sharp microelectrodes (150–250 MΩ) with an amplifier operating in bridge mode, into individual amacrine cells. Current and the visual stimulus were aligned in time within 0.1 ms, to allow reproducible recordings across the same visual and current stimuli presentations. The current amplitudes were chosen so they maintain the membrane potential within a physiological range. The estimated membrane resistance measured by using pulses of current was 40 ± 17 MΩ, and the measured membrane time constant was 17 ± 7 ms (n = 6). The current filtered to a 0–50 Hz bandwidth was convolved with an exponential filter of the appropriate amplitude to estimate the resulting membrane potential standard deviation (9.6 ± 3.63 mV)^22^.

### Measuring linear-nonlinear cascade models of the recorded responses

Recordings were used to produce LN models of visual responses and of responses to current injection^43^. The input to the model could be a visual stimulus or a current stimulus, and the output of the model could be either the predicted ganglion cell’s firing rate or the predicted amacrine cell’s membrane voltage. The estimated linear filters ranged from 200 to 600 ms in duration, and were estimated using white noise stimuli of length 300-1200 s.

To estimate the parameters of the LN model we used a semiparametric approach, in which the filter *f*(*t*) and the nonlinearity *N*(*g*) were computed at once. For a Gaussian white noise input, the optimal linear filter is indeed the Wiener system solution, which was computed in the Fourier domain by equation (1)^44,96^.

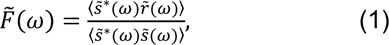

where 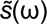 is the Fourier transform of the stimulus *s*(*t*), 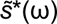 its complex conjugate, 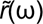 is the Fourier transform of the response *r*(*t*), and 〈…〉 denotes averaging over 1 s segments spaced every 0.1 s throughout the recording. The stimulus intensity *s*(*t*) was adjusted to have a zero mean. The denominator is the input auto-correlation, which corrects for possible deviation from the white Gaussian solution due to possible correlation structures in the input. For fitting of the ganglion cell spiking response, spike trains were converted into a continuous firing rate by binning the spike times in 1 ms intervals.

The predicted linear ganglion cell’s firing rate response or the amacrine cell’s membrane voltage *g*(*t*) was computed by convolving *f*(*t*) with the stimulus *s*(*t*).

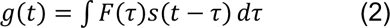

Then, a nonlinear function *N*(*g*) mapping the response to the linear prediction *g*(*t*) was found by computing the average value of *r*(*t*) over bins of *g*(*t*) containing an equal number of points. The output of *N*(*g*) determines the instantaneous firing rate^97^.

The amplitude of the filter was normalized such that the variance of the filtered stimulus, *g*(*t*), was equal to the variance of the stimulus, *s*(*t*) (equation (3)), to avoid the ambiguity existing in the scaling of linear filter y-axis and the nonlinearity x-axis.

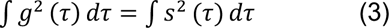

With this normalization, *f*(*t*) summarizes temporal processing, and *N*(*g*) captures the sensitivity of the stimulus.

Finally, the prediction of the LN model was calculated as

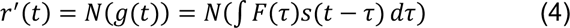

Spikes are then assumed to be generated by a Poisson process with a rate equal to *N*(*g*).

### Computation of amacrine pathway filter or feature

Amacrine visual response LN models were constructed using a white noise visual stimulus with a Gaussian distribution and the amacrine’s membrane potential recorded intracellularly (Fig. S1A). Similarly, a linear filter corresponding to amacrine cell transmission was computed by correlating the white noise Gaussian current signal with the spiking response of the ganglion cell, and then a static nonlinearity was computed by comparing the current linear prediction and the firing rate of the ganglion cell. These linear and nonlinear stages together were called the amacrine cell’s transmission LN model (Fig. S1B). Since the injected current and the visual stimulus were uncorrelated, the visual stimulus could be excluded in computing the amacrine transmission filter.

The output of the first LN model was the amacrine cell’s membrane voltage, which has already passed through the filtering of the amacrine cell’s membrane time constant. However, the input to the second LN model was the white noise Gaussian current injected to the amacrine cell, which is also filtered through the amacrine cell’s membrane time constant before inducing a voltage across the amacrine cell’s membrane. To compensate for this double counting of the membrane time constant, we deconvolved the transmission filter computed by reverse correlation by the amacrine’s membrane time constant (equation (5))^22^. The membrane time constant of the cell was measured by applying current pulses of 1 s. An exponential filter (equation (5)) was fit to the voltage response with membrane time constant τ.

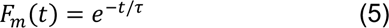

### Computing the output of amacrine pathway

Once the complete amacrine pathway model was obtained, the output signal of the amacrine pathway was computed during the current injection, which was used to study the effect of the amacrine pathway on the response of the ganglion cells. During current injection, the amacrine membrane voltage was produced by two sources of currents including the white noise Gaussian current injected to the amacrine cell and the current generated by the visual stimulation of the amacrine cell. We assumed for these two currents to be linearly combined in the amacrine cell before transmission to the ganglion cell, where the magnitude of their individual contributions (linear weights) was obtained by an additive least-squares model. This summed current was then convolved with the voltage transmission filter and then passed through the transmission nonlinearity.

### STA and STC analysis

The STA represents the difference between the mean (center of mass) of the spike-triggered ensemble and the mean of the raw stimulus ensemble and is estimated as follows assuming that the raw stimuli have zero mean:

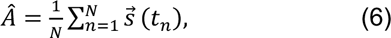

where *t*_2_ is the time of the *n*^th^ spike, 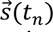 is a vector representing the stimuli preceding *t*_2_, and *N* is the total number of spikes. The stimulus 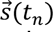 consisted of 60 stimulus values preceding a spike at a frame rate of 63 Hz and extending back in time 960 ms.

For a neuron with a single linear filter, the STA provides an unbiased estimate of the neuron’s linear filter (as obtained by reverse correlation in equation (1)), provided that the input has a spherically symmetric distribution, like a white Gaussian stimulus^60,96–98^, and the nonlinearity of the model is such that it leads to a shift in the mean of the spike-triggered ensemble relative to the raw stimulus ensemble.

To capture multiple features, the spike-triggered covariance (STC) technique was used^62,98^, where additional filters were recovered by seeking directions in the stimulus space in which the variance of the spike-triggered ensemble differs from that of the raw ensemble. Unlike the change in the mean that defines a single dimension, the variance can recover many dimensions at once.

The covariance matrix is formed from the outer product of the stimulus histories averaged across all spikes as follows.

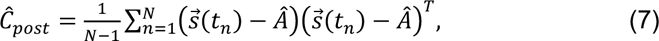

where *t*_2_ is the time of the *n*^th^ spike, 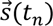 is a vector representing the stimuli preceding *t*_2_, and *N* is the total number of spikes, and *Â* is the STA (as computed in equation (6)).

*Ĉ_post_* contains the multidimensional variance structure of the spike-triggered stimulus ensemble, such that the variance of the STE along any direction *u* (*u* is a unit vector) is *u^T^Ĉ_post_ u*. We were interested in finding the features in the stimulus space for which the variance changes from that of the prior distribution of stimuli, which is Gaussian along all stimulus dimensions, independent of the response. We first construct the covariance difference matrix by subtracting *Ĉ_prior_* from *Ĉ_post_*.

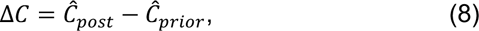

where 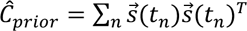.

Then the posterior second moment matrix *Ĉ_post_* can in general only differ from *Ĉ_prior_* in directions spanned by the model filters^49^. An eigenvector analysis of this matrix can determine the stimulus directions that account for most of the variance.

### Computation of orthogonal features

To model the other parts of the circuit feeding into the ganglion cell, we focused on the subset of stimuli that were not encoded by the amacrine cell pathway. This procedure isolates the subspace of the visual stimulus that has zero output through the amacrine cell pathway, called the null space of the amacrine pathway feature^46,47^. To do this, we orthogonalized the stimulus space with respect to the direction of the total amacrine pathway feature using a Gram-Schmidt procedure. Then we computed the ganglion cell’s STA in this new stimulus space as follows,

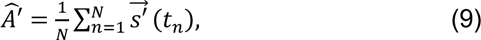

where 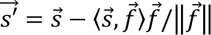, 〈…〉 denotes the inner products of two vector arguments, and 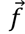 represents the amacrine pathway feature. We refer to *Â*′ as ganglion cell’s orthogonal STA (oSTA).

Similarly, we used the STC approach to recover the entire linear subspace representing the space spanned by the features encoded by the multiple pathways feeding into a ganglion cell that does not include the amacrine pathway feature. To find such features, we first projected out the amacrine pathway feature from the stimulus space, and constructed the sample second moment matrix of the spike-triggered ensemble in the new stimulus space and the difference covariance matrix. Then we found its eigenvectors and corresponding eigenvalues as explained above.

These eigenvectors identify the dimensions in the stimulus space along which the variance of spike-triggered stimulus distribution changes from that of the prior distribution of stimuli. Therefore, each eigenvector of this matrix represents a visual feature that might elicit spikes in the ganglion cell. The associated eigenvalue is equal to the variance of spike-triggered stimuli along this feature direction. Because the prior stimulus covariance matrix is subtracted from the posterior stimulus covariance matrix, an eigenvalue of zero corresponds to the prior variance and indicates that its associated eigenvector is not a relevant feature for ganglion cell response. Thus to find the relevant subspace, we extract the eigenvalues of the difference covariance matrix that are significantly different from zero. The number of significant eigenvalues gives an estimate for the dimensionality of the relevant stimulus space.

### Assessing the number of significant orthogonal ganglion cell features

Figure S5 shows the eigenvalue profile for a sample amacrine–ganglion cell pair, in which the eigenvalues of the covariance difference matrix were sorted from large to small values. The majority of the eigenvalues descend gradually indicating random fluctuations due to finite sampling (the white noise covariance difference matrix should have had constant eigenvalues at zero).

To assess the significance of eigenvalues we adopted a variety of significance tests, as sometimes the insufficient number of samples along some dimensions due to the low firing rate of the neuron, caused noisy estimations for the confidence intervals. In general, we preferred more qualitative definitions of significance, i.e., choose all eigenvectors corresponding to eigenvalues, which appear qualitatively different in magnitude from the “bulk spectrum” (Fig. S5). However, due to orthogonalization there were cases where we could detect eigenvalues that were significantly below or above the gradually descending region, but with arbitrary associated eigenvectors. For these cases, we used quantitative significance tests by computing the eigenvalue spectrum as a function of the number of spikes for a range of data fractions. Eigenvalues that were stable with respect to the number of spikes used in the analysis were judged to be significant (green trace in (Fig. S6A). We also used a nested hypotheses testing approach to determine the number of relevant dimensions corresponding to significant increases or decreases in variance^52^ (Fig. S6B). The significance level was chosen to be (p < 0.05). To get more accurate results, we only included ganglion cells from which we recorded at least 1200 spikes, being at least 20 spikes per temporal dimension for 60-dimensional linear temporal filters.

### Parameterization of amacrine nonlinear effects

Our goal was to find a model of modulatory functions of amacrine cells, which is consistent with a set of experimentally estimated correlations between the stimulus and neuron’s response, and at the same time imposes as little structure as possible for the nonlinear interactions among different pathways.

To this end, we estimated the ganglion nonlinear response function using a nonlinear regression framework defined as

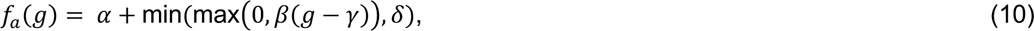

meaning that *f_a_*(*g*) is the linear function *α* + *β*(*g* − *γ*), except that it is limited to have a minimum value of *α* and maximum of *α* + *δ*. Here *α*, *β*, *γ*, and *δ* represent respectively the response baseline, sensitivity, threshold and gain, *g* represents the filtered stimulus by each ganglion cell’s visual feature identified using the STC analysis, and *f_a_*(·) denotes the ganglion cell firing rate response, which can vary as a function of the output of the amacrine pathway indicated by *a*. The estimated parameters *α*, *β*, *γ*, and *δ* change depending on the output of the amacrine pathway, which in turn depends on the specific stimulus sequence. Thus the set of parameters, *α*, *β*, *γ*, and *δ* is controlled by the amacrine pathway, and describe the immediate ganglion cell visual response to light. The parameters *α*, *β*, *γ*, and *δ* were found through a constrained nonlinear least-squares optimization. *α* and *β* were bounded to 0 from below and to the maximum value of the binned spiking probability of a ganglion cell representing its firing rate from above. *γ* was bounded between the minimum and maximum values of the filtered stimulus values for each ganglion cell’s feature, and *β* was unconstrained. Because the nonlinearities for some amacrine–ganglion cell’s feature pairs were strongly nonmonotonic, as a goodness-of-fit criterion, only the amacrine–ganglion cell’s feature pairs with adjusted r-square values greater than 0.7 (75^th^ percentile) across all four levels of amacrine pathway polarization were included in the analysis. Because the STC analysis gives orthogonal features, each ganglion cell feature may be created by multiple neural pathways, and thus each nonlinearity may reflect combined effects of the amacrine cell on those pathways.

The gain-sensitivity modulation (GSM) index was defined as the Pearson correlation coefficient between the amacrine pathway output and the ratio of gain to sensitivity, *δ*/|*β*|, where the parameters *δ* and *β* were taken from (equation 10) fit as above to nonlinearities at four different levels of amacrine output. Because amacrine cells were inhibitory, negative value of the GSM index show a greater reduction of ganglion cell gain than sensitivity, and positive values show greater reduction of sensitivity than gain.

### Consistency of amacrine effects with possible presynaptic or postsynaptic modulation

To assess whether modulatory control over different visual features associated with individual amacrine–ganglion cell pairs might act separately on specific visual features or act more similarly on all features, the pairwise Pearson correlation between vectors of effect types was computed. Each effect type vector represents the change in parameters of sensitivity, gain, and threshold for the fitted sigmoidal nonlinearities associated with each amacrine-ganglion cell pair falling in any of the two modulation classes. The amacrine-ganglion cell pairs for which at least three visual features with a nonlinear fit performance greater than 70% and the threshold and GSM modulation magnitudes greater than 0.2 were obtained, were included in this analysis. Note that some amacrine-ganglion cell pairs may appear in both classes, but distinct subsets of the visual features of the ganglion cell fall in each of the two classes. This suggests that individual amacrine cells can modulate the same ganglion cell using differential modulatory computations at multiple synaptic locations. The larger average correlation value suggests more similar effect type vectors across all the ganglion cell’s features for individual amacrine-ganglion cell pairs given a modulation class. The more similar effect type vectors indicates the greater similarity of the impact of a certain amacrine cell on multiple features of the same ganglion cell, and can be interpreted as a postsynaptic modulation by that amacrine cell. On the other hand, the less similar effect type vectors may imply that the impact from a particular amacrine cell is delivered to different presynaptic terminals that could have different nonlinear properties.

## Data and code availability

The datasets generated and analyzed in this study will be available on public repositories upon the acceptance of the paper.

## Acknowledgments

We thank K. Boahen, S. Ganguli and K. Shenoy for helpful discussions. This work was supported by grants from the NEI, Pew Charitable Trusts, McKnight Endowment Fund for Neuroscience, the Alfred P. Sloan Foundation and the E. Matilda Ziegler Foundation (S.A.B.).

## Author Contributions

N. N., M. M. and S.A.B. designed the study, N. N. and M. M. performed the experiments, N. N. performed the analysis, and N. N. and S.A.B. wrote the manuscript.

