## Supplementary information for "Modulation of input sensitivity and output gain by retinal amacrine cells"

**This PDF file includes:**

Figures S1 to S8

### Supplementary Figures

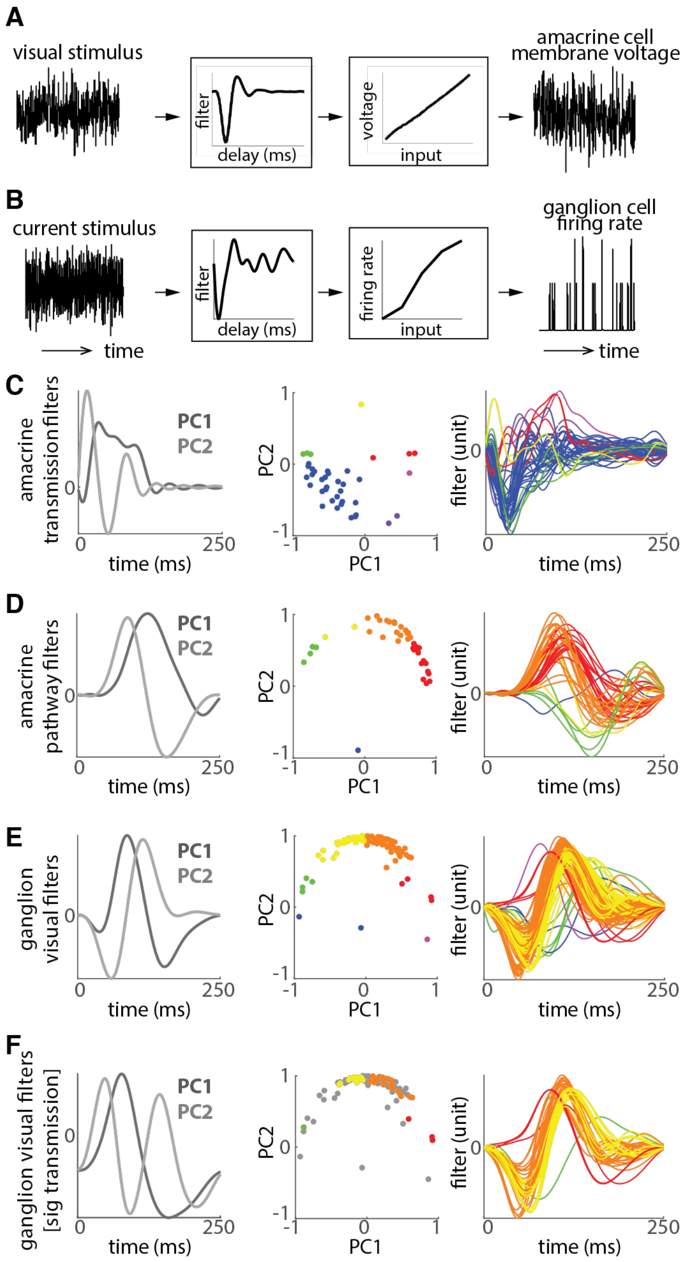


**Fig. S1. The visual feature of an amacrine pathway.** (A) The linear filter and nonlinearity between a visual stimulus and a sustained Off amacrine cell's membrane potential. This linear filter is referred to as the amacrine visual filter. (B) The transmission linear filter and nonlinearity between current injected into an amacrine cell and a ganglion cell firing rate response. The amacrine pathway filter is obtained by cascading the amacrine visual filter and the amacrine transmission filter after compensating for the amacrine cell membrane voltage time constant. (C) From left to right: the first two principal components of 39 significant amacrine transmission filters (accounting for 78% of the variance); projection of all amacrine transmission filters onto these two principal components (the points are coded by eight possible colors based on their angle around the origin of the two principal components plane); significant transmission filters color-coded according to their projection angle from the middle plot. (D) From left to right: the first two principal components of 39 amacrine pathway filters (accounting for 84% of the variance); projection of all amacrine pathway filters onto these two principal components; amacrine pathway filters color-coded according to their projection angle from the middle plot. (E) From left to right: the first two principal components of ganglion cell visual filters (accounting for 66% of the variance) based on responses of 104 recorded ganglion cells with firing rate greater than 0.5 Hz; projection of all ganglion cell filters onto these two principal components; visual filters of the ganglion cells color-coded according to their projection angle from the middle plot. (F) From left to right: For the subset of ganglion cells from (E) for which a significant transmission filter was measured between an amacrine–ganglion cell pair (referred to as affected ganglion cells), the first two principal components of ganglion cell visual filters (accounting for 74% of the variance); projection of all ganglion cell filters onto these two principal components, where colored points represent a subset of points in the projection scatter diagram in (E) corresponding to affected ganglion cells; and their corresponding ganglion visual filters.

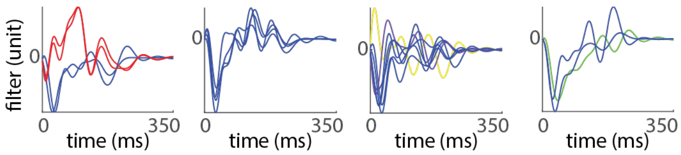


**Fig. S2. Diversity of amacrine transmission dynamics for single amacrine cells versus multiple ganglion cells.** Each plot shows the significant transmission filters from one amacrine cell to the affected ganglion cells simultaneously recorded using multielectrode arrays in each experiment session, for four sample amacrine cells. The colors represent the type of transmission filter as classified according to the principal component analysis in figure S1C.

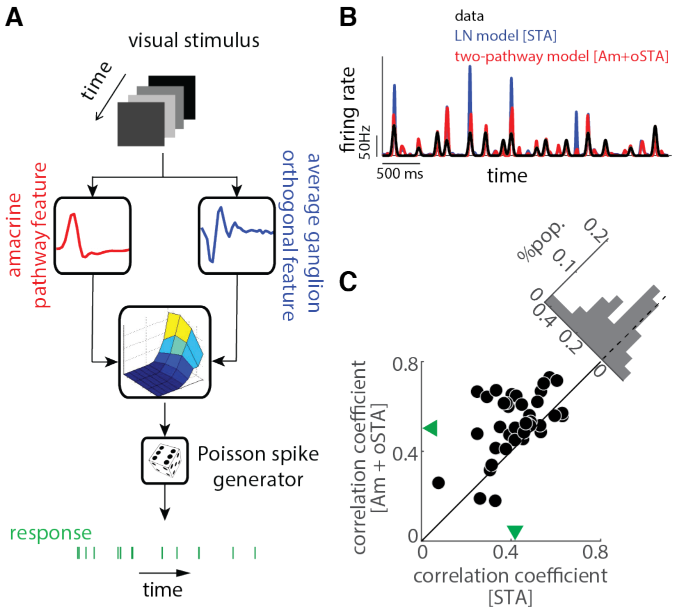


**Fig. S3. A model using the measured amacrine pathway more accurately predicts the ganglion cell response.** (A) Diagram of the two-pathway model used to predict the ganglion cell's firing rate response. The response of a ganglion cell was fit to the visual stimulus with a model containing two main pathways. The left pathway is the amacrine pathway visual feature (red filter), obtained by cascading the amacrine temporal receptive field and amacrine transmission kinetics. The right pathway (blue) is the average of other visual features (oSTA) encoded by the ganglion cell excluding the feature conveyed by the amacrine cell. Then, the outputs of these two pathways were combined through a two-dimensional instantaneous firing rate nonlinearity to predict the response of the ganglion cell (green spike train). (B) Illustrates the prediction of the model in (a). The actual firing rate (black) is plotted together with the predicted firing rate of a ganglion cell for an ~4-s stimulus segment from the test data, which was not used for fitting the model using an LN model with STA (blue) and using the two-pathway model with oSTA and the amacrine pathway visual feature (Am). (C) Shows the models' performance, measured as correlation coefficients between the ganglion cell firing rate and the predicted firing rate using the LN model [STA] (y-axis) versus the two-pathway model in (A) [Am+oSTA] (x-axis), for 39 ganglion cells with significant amacrine transmissions, over the withheld test data. Histogram in the upper right shows the population distribution of change in performance between the two models (0.10$\pm$0.13 mean$\pm$SD; $p = 1.65 \times 10^{-5}$ one-sided Wilcoxon signed-rank test). Arrowheads along the x- and y-axes mark the means of each distribution.

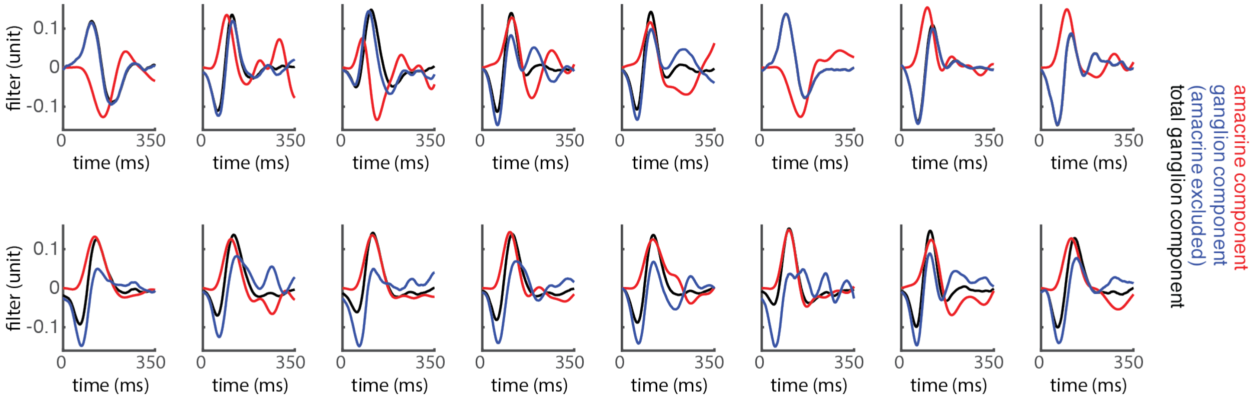


**Fig. S4. Relationship between the similarity of STA and oSTA and the relative temporal dynamics of amacrine-ganglion cell pairs.** Plots show the amacrine pathway feature (red), ganglion cell's STA feature (black), and ganglion cell's oSTA feature (blue), for sample amacrine-ganglion cell pairs. The samples are organized into two groups based on the distribution of the distance between STA and oSTA (Fig. 2C), corresponding to the 30$^{th}$ percentile of distance (top row), and the 70$^{th}$ percentile of distance (bottom row).

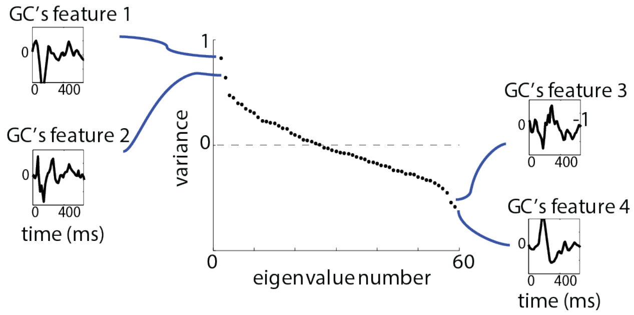


**Fig. S5. Eigenvalue profile of the STC matrix of the stimulus ensemble orthogonal to the amacrine pathway feature.** For this sample amacrine–ganglion cell pair (the same sample used in Fig. 3C), four significant STC dimensions corresponding to four significant eigenvalues were found. The total stimulus dimension was 31, corresponding to 31 16-ms uniform field stimulus frames changing their intensity randomly over time.

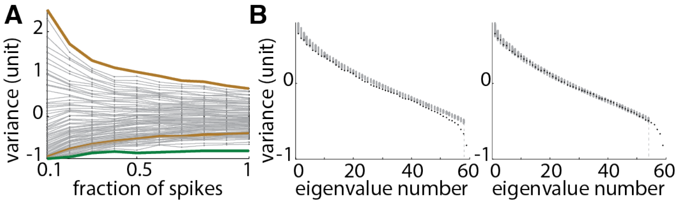


**Fig. S6. Significance test of STC features.** (A) The eigenvalue spectrum was computed for different fractions of data, ranging from one-tenth to the entire number of spikes. Those eigenvalues that remained stable across different fractions of data (sample green trace) are considered significant eigenvalues, i.e., those that are different from the bulk spectrum due to the spiking events and not due to random fluctuations along poorly sampled dimensions. Those eigenvalues that vanish (sample brown traces) when the size of data increases are considered nonsignificant. (B) A nested bootstrap was performed to test the significance of each eigenvalue. The null hypothesis starts with the assumption that there is no significant eigenvalue, then a confidence interval for each eigenvalue is computed, if at least one eigenvalue falls outside of the confidence interval the null hypothesis is rejected (left plot), and the next null hypothesis with the assumption of only one significant eigenvalue is tested, and this procedure continues until a null hypothesis is accepted (right plot), i.e. assuming the null hypothesis, no eigenvalue falls outside of the confidence interval. For the example shown here four significant eigenvalues were found.

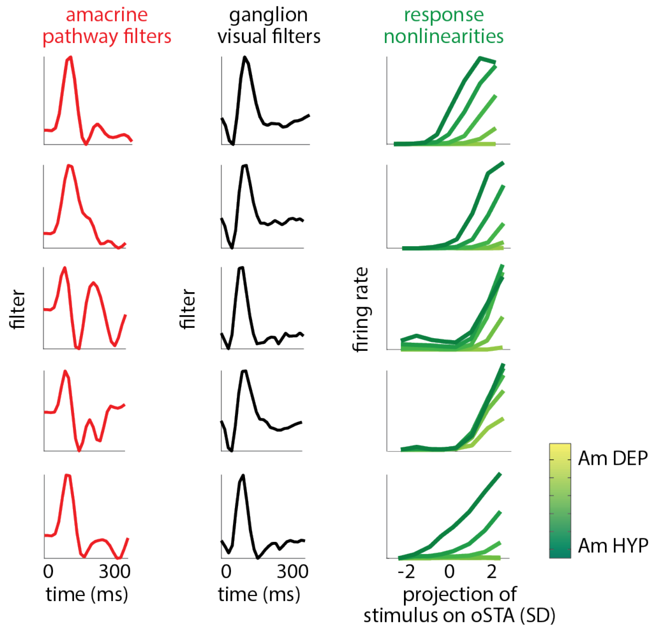


**Fig. S7. Diversity both in linear transmission and in nonlinear modulatory effects from one amacrine cell to multiple ganglion cells.** The plots illustrate the linear and nonlinear interactions between one sample amacrine cell versus five ganglion cells recorded simultaneously using multielectrode arrays. Each row represents a pair of amacrine and ganglion cells where the amacrine cell is the same across rows. Red traces (first column) represent amacrine pathway filters between the amacrine cell and each ganglion cell. Black traces (second column) represent ganglion cell visual filters for each of the five affected ganglion cells in the experiment. Green traces (third column) show ganglion cell firing rate as a function of the projection of the stimulus on the ganglion cell visual features with the amacrine pathway feature excluded, at five levels of amacrine cell polarization.

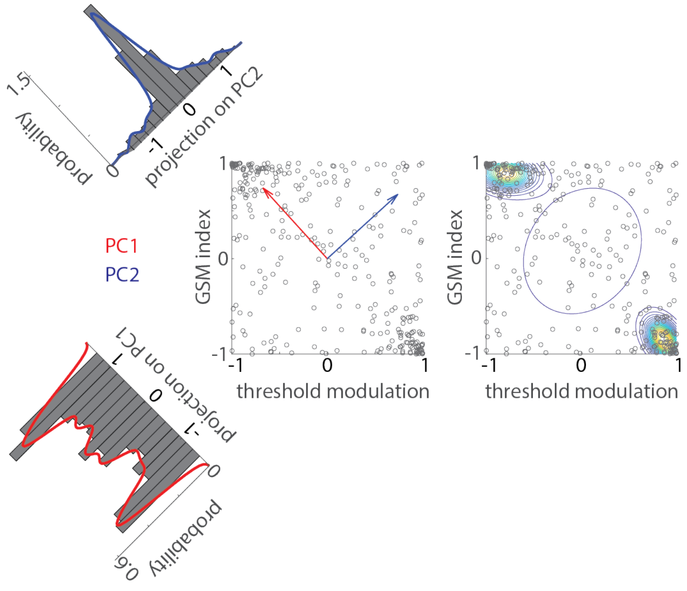


**Fig. S8. Cluster analysis of nonlinearity modulations.** The scatter plots show the distribution of the GSM index and threshold modulation values measured for all the 321 amacrine–ganglion cell's feature pairs (the data used for generating the two-dimensional histogram in figure 4D). The two arrows on the left scatter plot indicate the directions of the first (red) and second (blue) principal components of the data points. The diagonal histograms on the left show the distribution of the projected data points onto the first (bottom histogram) and second (top histogram) principal components. The red and blue curves over the histograms are the fitted probability density functions obtained using a kernel density estimator with a normal smoothing function, for the data points projected respectively onto the first and second principal components. Using a gaussian mixture model identified two main clusters representing two distinct classes in the space of nonlinearity modulations, as shown by two groups of contours at the corners of the right scatter plot. The colored ovals indicate the contours of the two-dimensional gaussian probability density functions fitted on the data points. The left and right scatter plots are the same.
